# Modulation of huntingtin S421 phosphorylation in a Huntington’s disease mouse model and its detection in nonhuman primate cerebrospinal fluid

**DOI:** 10.1101/2025.07.18.665500

**Authors:** Margherita Verani, Paola Martufi, Lara Petricca, Andrea Caricasole, Maria Carolina Spiezia, Olof Larsson, Licia Tomei, Valentina Fodale, Ignacio Munoz-Sanjuan, Thomas F. Vogt, Celia Dominguez, Elizabeth M. Doherty, Ramee Lee, Cristina Cariulo

**Affiliations:** Translational and Discovery Research Department, IRBM S.p.A. Via Pontina km 30.600, 00071 Pomezia, Rome, Italy; CHDI Management, Inc., the company that manages the scientific activities of CHDI Foundation, Inc., Princeton NJ 08540, USA

## Abstract

Huntington’s disease (HD) is a progressive neurodegenerative disease caused by the pathologic expansion of a CAG repeat in the first exon of the huntingtin (*HTT*) gene, resulting in a huntingtin (HTT) protein with an expanded polyglutamine (polyQ) tract. Phosphorylation at residue S421 (pS421) is one of the post-translational modifications proposed to influence the biology of wild-type and mutant (m)HTT, such as HTT stability and clearance, HTT subcellular localization, mHTT toxicity, and regulation of HTT function in axonal transport. However, the detection and quantification of S421-HTT phosphorylation in relevant biological contexts have remained challenging and the consequences of pS421 in HD pathogenesis remains unclear. Here we report the development of a novel ultrasensitive immunoassay enabling the specific and sensitive detection of pS421-HTT in a variety of biologically relevant contexts. With this assay we conducted a longitudinal assessment of pS421 levels in tissues from a mouse model of HD to investigate the relationship between S421 phosphorylation and phenotypic progression. We also identified PRKACA, the cAMP-regulated catalytic α subunit of PKA, as a kinase capable of phosphorylating S421-HTT, demonstrating its ability to regulate endogenous pS421 in human cells. Finally, we exploited the sensitivity of the assay to detect endogenous pS421-HTT in cerebrospinal fluid (CSF) from nonhuman primates, showing for the first time that phosphorylation at S421-HTT can be detected in this bio-fluid. These reagents and assay will enable investigation of the biological consequence and the relevance of pS421 in the natural history of HD.

## Introduction

Huntington’s disease (HD) is caused by a pathologic CAG expansion within the first exon of the huntingtin (*HTT*) gene which results in a mutant protein (mHTT) bearing an expanded polyglutamine (polyQ) repeat. Like in several other neurodegenerative diseases, HD is associated with misfolding and accumulation of this disease-related protein, and protein function is frequently linked to post-translational modification (PTM). Modifications such as phosphorylation can have profound implications for the structure-function and pathology associated with neurodegeneration in HD, thought to be the result of both gain and loss of function mechanisms [1–5]. Although the cryoEM-structure of full-length HTT complexed with its partner HAP40 has been solved [6], the relationship between HTT structure and its biological activity is still poorly understood, at least in part due to the pleiotropic functions and structural complexity of HTT, including several unstructured regions [3, 4, 6, 7].

Current research in the development of therapeutics for HD include a focus on lowering HTT levels in the CNS through different approaches [8] and, consequently, the capacity to accurately quantify HTT levels in HD models and in persons with HD (PwHD) has been critical [9, 10]. The demonstration that lowering HTT levels can provide benefit in HD animal models [11] has spurred interest in different approaches to modulate *HTT* transcription, HTT protein synthesis and clearance [8]. Generally, these approaches are focused on reducing *HTT* transcription or mRNA levels/translation (see for examples [11, 12]).

Phosphorylation of S421 (pS421; numbering based on polyQ 23) has been implicated as a modulator of HTT steady-state protein levels in cells and an HD mouse model, with a phosphomimetic S421D mutation resulting in amelioration of the disease phenotype [13]. pS421 has also been implicated in the regulation of several HTT functions that are impaired in HD, such as subcellular localization [7, 14], axonal transport [15–18], and mHTT toxicity [7, 13, 19–21]. To further evaluate the hypothesis that pS421-HTT is of pathophysiological, and perhaps even therapeutic, relevance requires a robust means to detect and quantify phosphorylation to enable identification of modulators for proof-of-concept studies in preclinical HD models. It has been reported that pS421-HTT can be detected in cells and brain tissue by mass spectrometry [7, 22, 23] as well as by western blotting employing polyclonal antibodies (pAbs; [19, 24]). However, there are currently no quantitative and sensitive methods for pS421-HTT detection to enable the identification of candidate kinases and small-molecule modulators for the profiling of this PTM in HD models.

We previously reported an ultrasensitive, quantitative, sandwich immunoassay to study phosphorylation of T3-HTT [25] and S13-HTT [26] that utilized single molecule counting (SMC) technology. This approach has been applied to detect HTT in human cerebrospinal fluid (CSF) in clinical trials [9]. Here we report the use of this technology to assess pS421-HTT, further refined by the development of a new monoclonal antibody specifically characterized for this application. We assessed the relevance of pS421-HTT in a longitudinal study in a mouse model of HD, showing a distinct decrease in striatal and hippocampal pS421 correlated to phenotypic progression. We also report the identification and validation in a cell based of PRKACA as a kinase able to phosphorylate S421-HTT, providing a valuable tool to further investigate pS421-HTT biology. Additionally, the sensitivity of our assay allowed us, for the first time, to investigate HTT phosphorylation in cynomolgus macaque CSF as a proof-of-concept for using this assay in human CSF samples. Overall, our assay provides a means to assess the potential role of this PTM in biomarker studies of HD [25].

## Results

### Development of an SMC assay for pS421-HTT

To develop an assay to detect and quantify pS421-HTT with high sensitivity in a variety of relevant biological matrices/contexts, we utilized the SMC platform to implement a similar strategy to our previously published HTT pT3 and pS13 immunoassays [25, 26]. We initially employed two companion SMC antibody pairings, where the same capture antibody (MW1) was coupled with two distinct detection antibodies: mAb 2B7 for the detection of HTT protein (regardless of phosphorylation state), and polyclonal antibody (pAb) Ab2174 for detection of pS421-HTT. The latter is a commercially available pAb (Abcam) that we used for our initial proof-of-concept studies and has a known rabbit-immunization peptide sequence (RSRSGpSIVE).

We verified the ability of Ab2174 to specifically recognize bona-fide phosphorylation at S421 by western blotting and the SMC assay on recombinant full length Q23 HTT proteins purified from HEK293 cells, with or without the phosphor-abrogative (S421A) and mimetic (S421D) mutations (Fig. 1A). We further validated Ab2174-based pS421-HTT readouts in a more complex biological context where HEK293T cells were transiently transfected with full length Q23 and Q48 HTT constructs, in the presence or absence of the S421A mutation. Ab2174 specifically detected pS421-HTT in both western blotting and the SMC assay, with a robust decrease in signal in samples overexpressing the corresponding S421A mutants (Fig 1B). A faint signal in these full length HTT S421A overexpressing and recombinant S421A and S421D sample proteins was observed, indicating a low level of reactivity of the commercial Ab2174 to these mutants.

**Figure 1.**
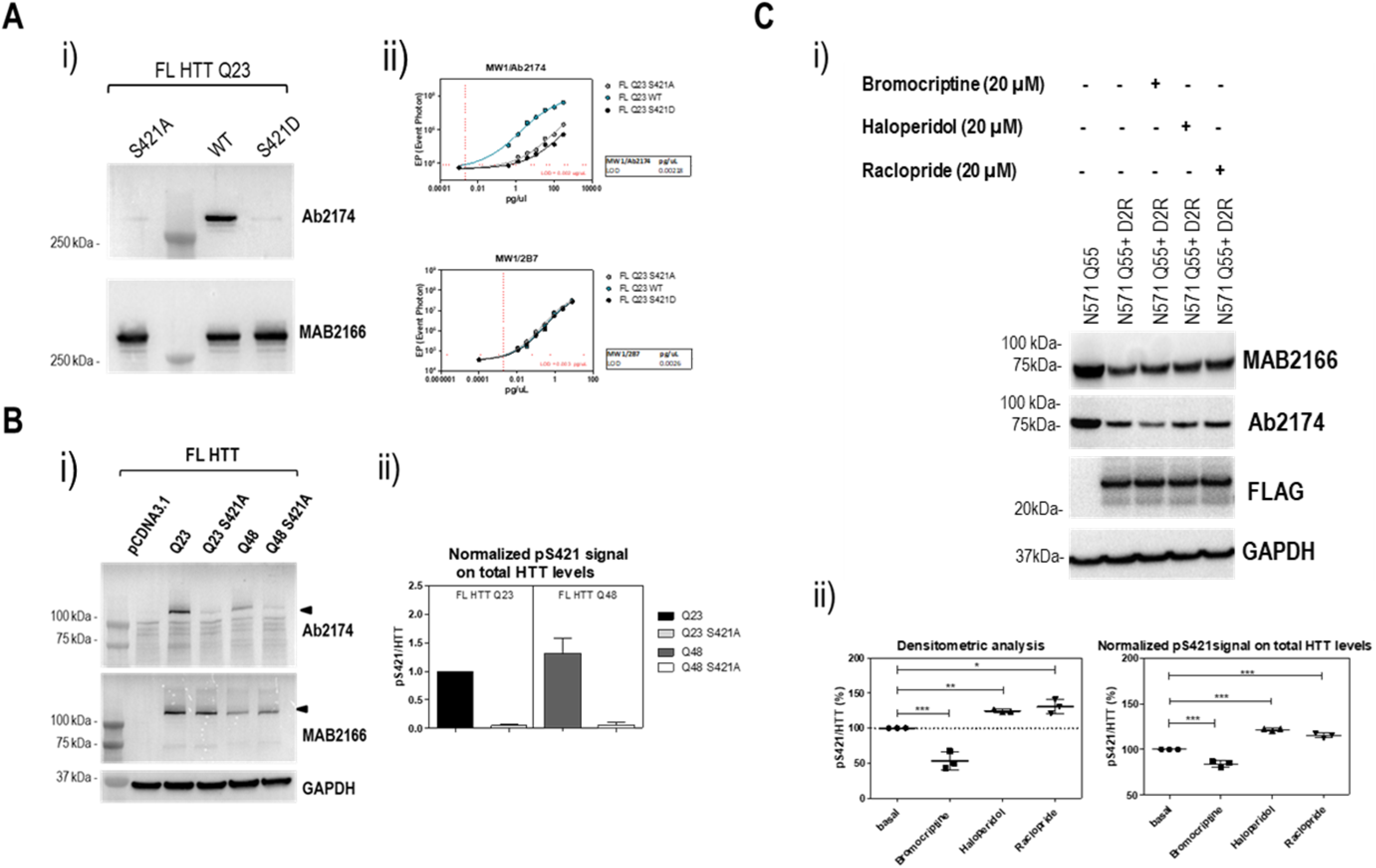
Development and validation of an SMC assay for the detection of pS421-HTT using the MW1-Ab2174 antibody pair. **A. Specificity assessment of Ab2174 for pS421 HTT by WB and the SMC assay in purified, recombinant FLQ23 HTT proteins with/without S421 A or D mutations**. i) WB probed with Ab2174 polyclonal antibody or MAB2166 antibody reveals pS421-HTT and total HTT levels, respectively. ii) SMC analysis of pS421-HTT levels (MW1-Ab2174) or total HTT levels (MW1-2B7) performed on serial dilutions of recombinant proteins in i). Representative experiment of n=3 is shown. **B. Specificity assessment of Ab2174 for pS421 HTT by WB and SMC assay in HEK293T cells overexpressing FL HTT with/without S421 A or D mutations.** i) WB probed with Ab2174 polyclonal antibody or MAB2166 antibody to demonstrate the levels of pS421-HTT and total HTT, respectively. Anti-GAPDH was used as loading control. ii) Normalized pS421 signal (MW1-Ab2174 SMC assay) on total HTT levels (MW1-2B7 SMC assay) performed on samples in i). Means and standard deviations from n=3 biological replica. **C. Pharmacological modulation of pS421-HTT levels in HEK293T cells transfected with N571 Q55 HTT protein and FLAG-tagged dopamine D2 receptor, treated with D2R agonist (Bromocriptine) or antagonists (Haloperidol, Raclopride).** i) WB probed with MAB2166 antibody, Ab2174 polyclonal antibody or FLAG antibody demonstrating total HTT levels, pS421-HTT and the D2 receptor, respectively. Anti-GAPDH used as loading control. ii) Densitometric analysis of WB in i) and normalized pS421 signal (MW1-Ab2174 SMC assay) on total HTT levels (MW1-2B7 SMC assay) performed on the same samples. Means and standard deviations from n=3 biological replica. Statistical analysis by one-way analysis of variance (*P<0.05; **P < 0.01; ***P < 0.005).

We then evaluated the SMC assay’s capacity to monitor pharmacological modulation of pS421-HTT levels. It was previously reported that pharmacological intervention of the dopamine D2 receptor (D2R) modulates pS421-HTT levels in HEK293T cells [27]; we therefore used the same conditions from this previous report to co-transfect HEK293T cells with D2R and 1-571-HTT constructs and incubated for 24 hrs with the D2R agonist bromocriptine or D2R antagonists haloperidol or raclopride (all at 20 µM). As shown by western blotting and the SMC assay (Fig. 1C), activation of D2R by the agonist bromocriptine resulted in a modest decrease in pS421-HTT levels, while the D2R antagonists produced a slight increase in pS421-HTT levels, with both direction and the modesty in magnitude of effect in agreement with previous reports [27].

pAbs have been used in the detection of HTT PTMs [19, 24, 28, 29]. mAbs are typically preferred for immunoassay development because they can be defined molecularly and expressed to allow unlimited homogeneous production [30]. To further optimize our assays, we developed a novel rabbit mAb following a stringent workflow. This comprised analysis of immunized rabbit antisera, B cell clone supernatants and purified IgGs for specific detection of S421 under both native (SMC assay) and denaturing (western blotting) conditions, resulting in the identification of candidate mAb (7D9) producing B-cells. The specificity of mAb 7D9 in the SMC assay was initially evaluated using recombinant HTT proteins and overexpressed HTT in HEK293T cells, analogously to the above procedure validating the pAb Ab2174.

In addition, although the mAb MW1 proved to be a functional capture antibody in the proof-of-concept studies, we wanted to avoid potential mAb MW1 bias for long polyQ expansions as previously shown [31]. We therefore replaced mAb MW1 with the mAb 2B7 whose epitope falls within the N-terminal region of HTT, thereby avoiding the polyQ stretch of Exon1. The rest of the experiments in this study therefore utilized 2B7 in combination with the commercially available mAb D7F7 (Cell Signaling Technology) as detection antibody to assess total HTT levels in the companion SMC assay to allow the same capture antibody for detection of pS421 [32].

We first validated the performance of the refined SMC assay, employing the 2B7-7D9 pair, on recombinant full length HTT proteins, with or without the phosphoabrogative S421A and phosphomimetic S421D mutations, using western blotting as an orthogonal readout (Fig. 2A). The same validation approach was done in HEK293T cells overexpressing full length HTT with or without the S421A mutation (Fig. 2B), confirming the high specificity of mAb 7D9 for pS421 also in biologically more relevant contexts, under both native and denaturing conditions.

**Figure 2.**
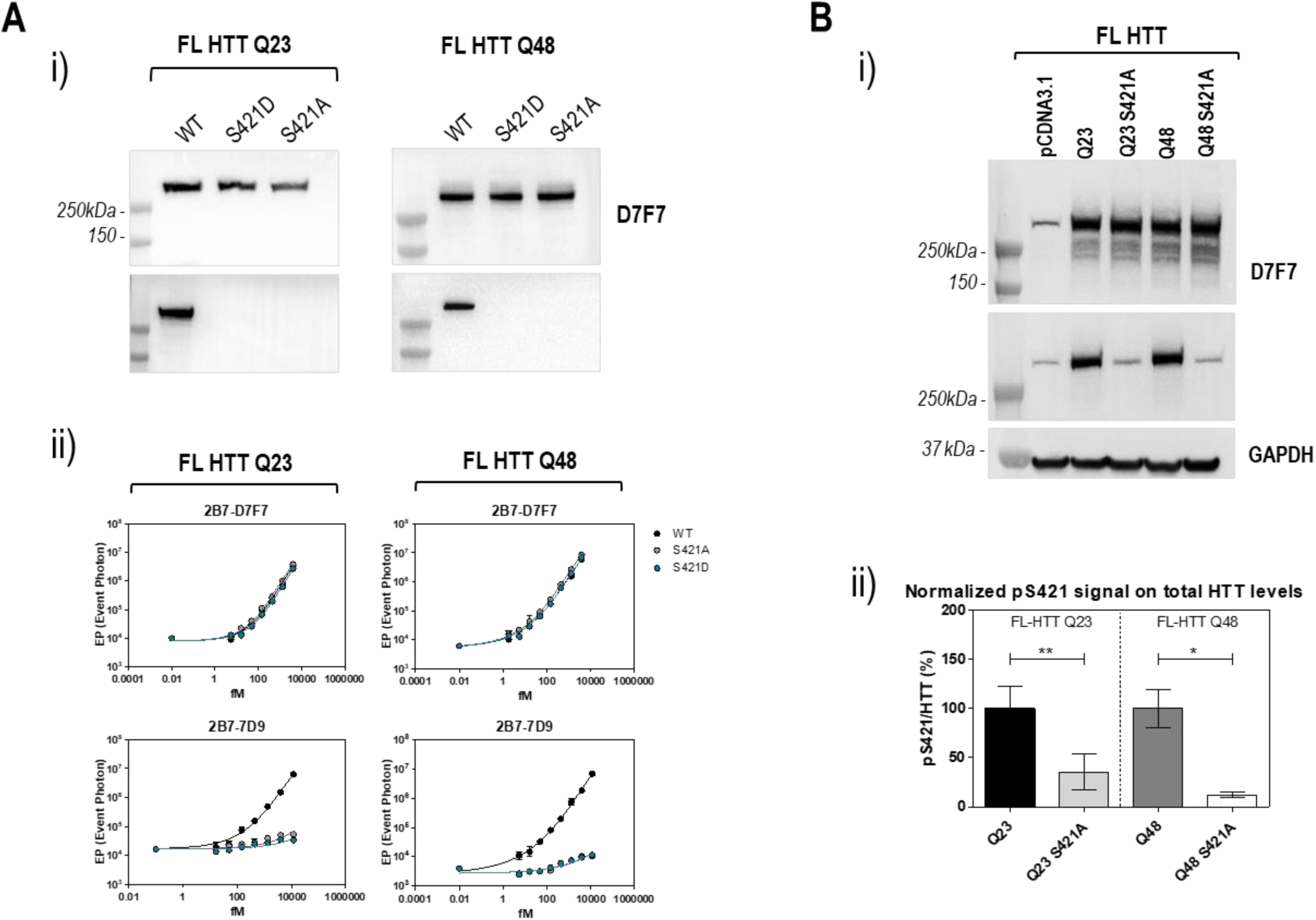
Optimization of the SMC assay to detect pS421-HTT by employing the 2B7-7D9 pair. **A. Specificity assessment of 7D9 for pS421-HTT by WB and SMC assay in purified, recombinant FLQ23 HTT proteins with/without S421 A or D mutations**. i) WB probed with D7F7 antibody or 7D9 monoclonal antibody revealed total HTT levels or pS421-HTT, respectively. ii) SMC analysis of total HTT levels (2B7-D7F7) or pS421-HTT levels (2B7-7D9) performed on the serial dilutions of recombinant proteins in i). **B. Specificity assessment of 7D9 for pS421-HTT by WB and SMC assay in HEK293T cells overexpressing FL HTT with/without S421 A or D mutations.** i) WB probed with D7F7 antibody or 7D9 monoclonal antibody determining total HTT or pS421-HTT levels, respectively. Anti-GAPDH used as loading control. Representative experiment of n=3 biological replica is shown. ii) Normalized pS421 signal (2B7-7D9 SMC assay) on total HTT levels (2B7-D7F7 SMC assay) performed on samples in i). Means and standard deviations from n=3 biological replica. Statistical analysis by one-way analysis of variance (*P<0.05; **P < 0.01).

To exclude that the decrease in signal obtained in the presence of the phosphomimetic or abrogative mutations was simply due to the modification of the HTT epitope rather than to the actual absence of phosphorylation at S421, we performed a dephosphorylation treatment on recombinant full length HTT (Q23 and Q48) and overexpressed full length Q48 HTT in HEK293T cells with or without S421A mutation. In both cases a virtually complete abolishment of pS421-HTT signal was found (Fig. S2), thereby confirming the capability of this pS421 mAb to recognize bona-fide phosphorylation.

Since HTT phosphorylation at S421 is well established to occur in HEK293T cells [7, 19, 23], the potential of mAb 7D9 to detect endogenous pS421-HTT was examined in this cell line by both western blotting and the SMC assay, and its specificity confirmed by immunodepletion (ID) of HTT. Immunoprecipitation of endogenous HTT in HEK293T cells with the specific HTT antibody (D7F7) produced a robust decrease in HTT levels in the ID samples compared to immunoprecipitation performed with an unrelated antibody (mock), as observed in both western blotting (probed with MAB2166 HTT antibody) and the SMC assay assessing total HTT levels (2B7-MAB2166) (Fig. 3A). To obtain a reliable orthogonal confirmation of the actual enrichment/depletion of HTT protein in the samples, we employed MAB2166 instead of mAb D7F7 because the latter was already used in the immunoprecipitation step. In parallel, mAb 7D9 antibody was shown to specifically detect endogenous pS421-HTT in HEK293T cells, as evident from the loss in signal in the ID samples as evaluated by western blotting and the SMC assay. The same approach was carried out in cortical tissues from *Hdh* Q7/Q175 mice, where endogenous HTT was pulled down with mAb D7F7 or mock antibody and the expected signal decrease in the immunodepleted samples of both total HTT and pS421-HTT was demonstrated by MAB2166 and mAb 7D9, respectively (Fig. 3B), confirming a high assay specificity to detect endogenous HTT in HD mouse model tissue.

**Figure 3.**
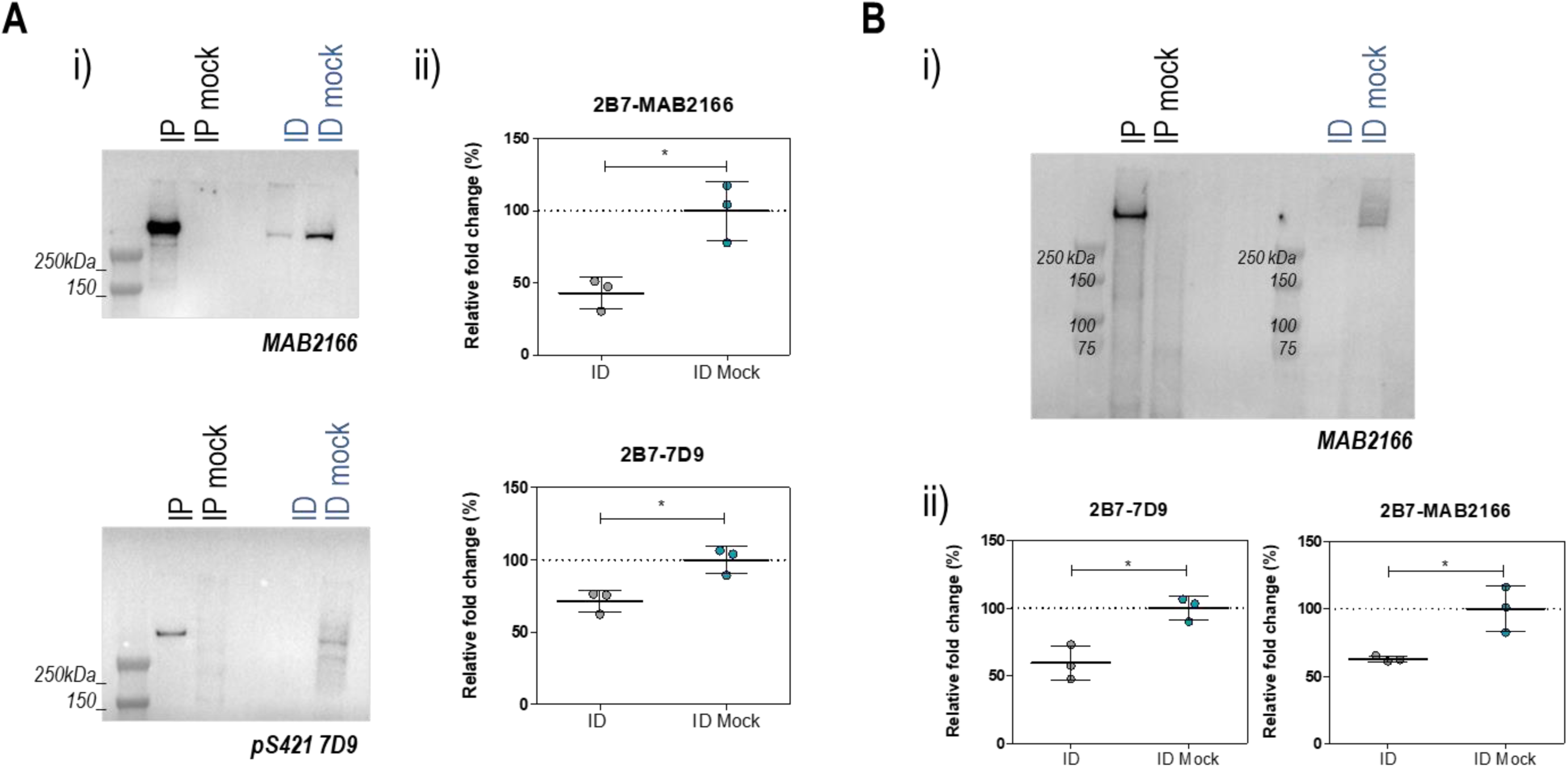
Endogenous pS421-HTT detection through the new 2B7-7D9 SMC assay. **A. Specificity of endogenous pS421-HTT detection in HEK293T cells.** i) Endogenous HTT from HEK293T cell lysate was pulled down with D7F7 or a mock antibody and analyzed by WB using MAB2166 or 7D9 monoclonal antibodies for total HTT and pS421 HTT levels, respectively. The presence or absence of HTT and pS421-HTT was assessed in immunoprecipitated samples (IP) and immunodepleted samples (ID), respectively. ii) SMC assay for total HTT levels (2B7-MAB2166) and pS421-HTT levels (2B7-7D9) performed on the ID samples in i). **B. Specificity of endogenous pS421-HTT detection in cortical tissue from *Hdh* Q7/Q175 mice.** i) Endogenous HTT from HD mice cortex homogenates was pulled down with D7F7 or a mock antibody and analyzed by WB probed with MAB2166 for total HTT levels. The presence or absence of HTT was verified in the IP and ID samples, respectively. ii) SMC assay for total HTT levels (2B7-MAB2166) and pS421-HTT levels (2B7-7D9) performed on the ID samples in i). Means and standard deviations of n=3 independent experiments. Statistical analysis by paired two-tailed *t*-test, (*P<0.05).

### HTT-S421 phosphorylation is reduced in CNS tissues from an HD mouse model

The ability to quantify pS421-HTT levels in cells and mouse tissues enabled the analysis of potential effects of polyQ expansion on S421-HTT phosphorylation. Previous studies have shown that phosphorylation at S421 is decreased in transgenic mice expressing full length mutant HTT [24]. However, lack of a method sensitive enough to quantify endogenous pS421 has precluded the conduct of a longitudinal study in preclinical HD models in sufficient detail to delineate the relevance of this PTM to HD pathophysiology and phenotypic progression.

We therefore set out to investigate pS421-HTT levels in CNS and peripheral tissues from *Hdh* Q7/Q7 and Q7/Q175 mice at different ages as a model of disease progression. Analysis of SMC data on pS421-HTT levels in all CNS tissues studied (cortex, hippocampus, cerebellum and striatum) from Q7/Q7 and Q7/Q175 *Hdh* mice at 2 months of age revealed a significant hypophosphorylation in HD mice compared to WT (Fig. 4Ai). Interestingly, a corresponding reduction in pS421 was not detected in heart tissue, pointing to a different modulation or underlying mechanisms leading to hypophosphorylation in the CNS versus peripheral tissue. The corresponding analysis of the same tissues from control versus HD model mice at 6 and 10 months of age further extended our findings that only brain regions exhibited a significant decrease in phosphorylation associated with polyQ expansion (Fig. 4Aii, 4Aiii). Of particular interest to the present study is the correlation between the degree of pS421-HTT modulation and HD progression. The striatal region of the CNS is known to be highly susceptible to neurodegeneration in HD [33–37], and indeed the hypophosphorylation effect evaluated here is more pronounced in this region, as well as in the hippocampus. Interestingly, increasing evidence suggested that extra-striatal dysfunction can make key contributions to the non-motor symptoms of HD [38, 39], with a specific focus on the HD-associated pathophysiology in the hippocampus [40]. Plotting hippocampal and striatal pS421-HTT levels in the Q7/Q175 mice over time confirmed the relative sensitivity of these two brain regions to phosphorylation with age, and the correlation of pS421-HTT status to disease pathogenesis (Fig. 4B).

**Figure 4.**
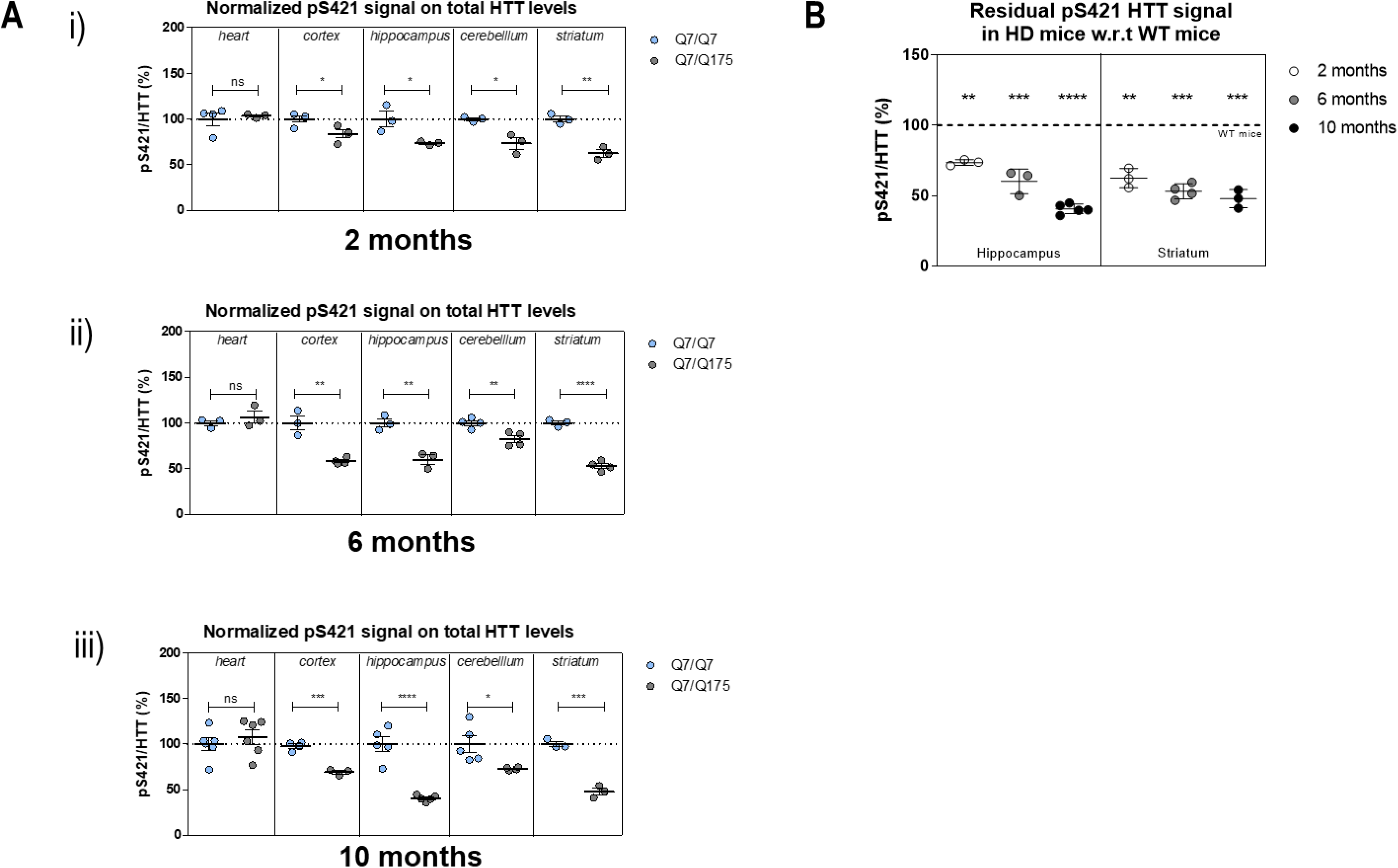
Decrease in endogenous pS421-HTT in CNS tissues from Q7/Q175 HD mice. **A. Determination of pS421-HTT in central and peripheral tissues from Q7/Q7 and Q7/Q175 mice at 2 and 6 months of age.** i) Normalized pS421 signal (2B7-7D9 SMC assay) on total HTT levels (2B7-D7F7 SMC assay) performed on a minimum of n=3 mice per genotype at 2 months of age. ii) Same assessment as in i) at 6 months of ages. Statistical analysis by unpaired *t*-test two-tailed (*P<0.05; **P < 0.01; ****P<0.0001). **B. Residual pS421-HTT levels measured in HD Q7/Q175 mice relatively to WT Q7/Q7 mice in striatal tissue at 2, 6 and 10 months of age**. Normalized pS421 signal (2B7-7D9 SMC assay) with total HTT levels (2B7-D7F7 SMC assay) performed on a minimum of n=3 mice per genotype expressed as percentage of the mean from WT Q7/Q7 mice, arbitrarily set to 100%. Statistical analysis by one-way analysis of variance, Tukey’s Multiple Comparison Test (*P<0.05) and Dunnett’s Multiple Comparison Test (***P<0.0001).

### PRKACA kinase is able to selectively modulate endogenous HTT-S421 phosphorylation levels in human cells

Next we sought to identify kinases capable of modulating endogenous pS421-HTT levels in cells. We profiled a collection of 370 full length kinases for *in vitro* phosphorylation of each of the four substrate peptides (Fig. S3A), based on the HTT sequence KEESGGRSRSGS_421_IVELIAGGG bearing S/A substitutions at different positions in the sequence. The resulting top 10 pS421-selective kinases (Fig. S3B) included the previously identified kinases AKT and SGK [19, 41], confirming our kinase profiling approach. To demonstrate their capability to modulate endogenous pS421-HTT levels, we transiently transfected HEK293T cells with expression plasmids encoding each of the identified top 10 kinases (SGK1-3, AKT1, MSK1, PRKACB, PRKACA, STK38L, PRKACG and PKN3) and examined the levels of pS421-HTT 48 hrs post-transfection by both western blotting and SMC assay. PRKACA was shown to be the only kinase capable of increasing the level of pS421 protein expressed from the endogenous *HTT* locus as measured using mAb 7D9 in western blotting (Fig. 5A and Fig. S4) and SMC assay (Fig. 5B).

**Figure 5.**
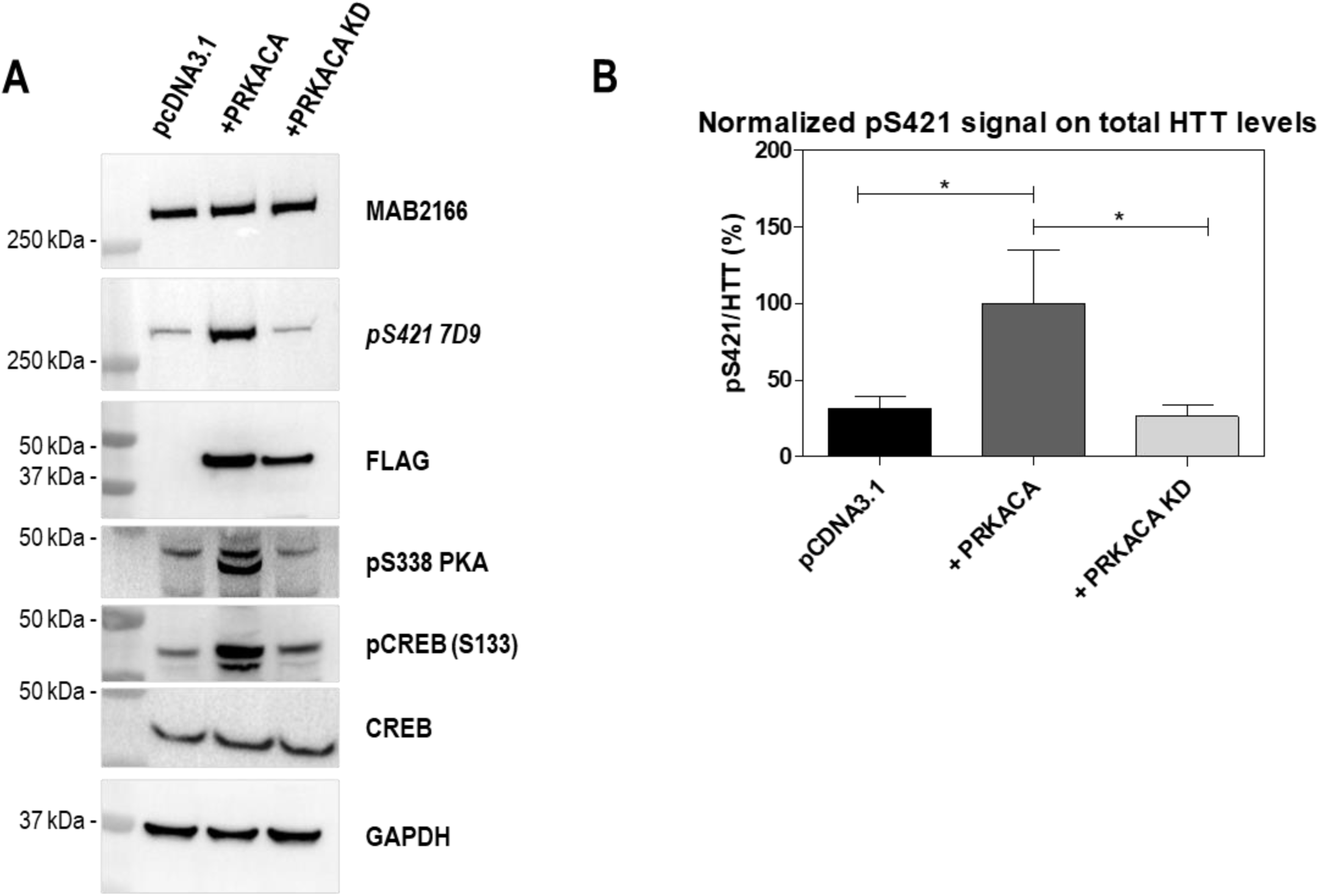
Identification of a tool kinase for pS421 HTT. **A. WB of lysates from HEK293T cells transfected with an expression plasmid for FLAG-tagged PRKACA or its catalytically inactive mutant.** Expression levels of endogenous HTT and pS421-HTT as detected by MAB2166 and 7D9, respectively. Immunoblotting with an anti-FLAG and anti-GAPDH were performed to control for correct kinase expression and as a loading control. WB probed with pS338 PKA, pCREB and CREB to indicate the activity of PRKACA kinase. Representative experiment of n=3 biological replica. **B. SMC data on pS421 (2B7-7D9) normalized to total HTT levels (2B7-D7F7) of the same samples in A.** Means and standard deviations of n=3 biological replica. Statistical analysis by one-way analysis of variance (*P<0.05).

It should be noted that neither AKT [19] nor SGK [41] has been shown to selectively increase endogenous pS421 HTT. The activity of PRKACA on pS421-HTT levels was dependent on the catalytic activity of the kinase, due to lack of effect using a kinase-dead mutant, defined as K/H substitution at residue 72 and G/V at residue186, which are known to affect both PRKACA autophosphorylation (S388;[42]) as well as phosphorylation of the bona-fide substrate CREB [43] (Fig. 5A). Furthermore, PRKACA did not significantly modulate phosphorylation at pT3 or pS13 (Fig. S5), two sites of HTT phosphorylation of functional relevance [25, 26, 28, 44–48], further demonstrating the specificity and selectivity of PRKACA to specifically phosphorylate the HTT protein at S421, at least relative to these two relevant HTT phosphorylation sites.

### Detection of phosphorylated S421-HTT in CSF from cynomolgus macaque

Identifying relevant disease biomarkers that could also be employed in a clinical setting would be a significant advancement for future clinical use in HD. The challenge in assessing protein levels in CSF lies in the high variability in CSF protein composition and a generally low protein abundance, with a particular difficulty assessing levels of protein phosphorylation. We therefore investigated the possibility of assessing levels of pS421-HTT in this highly relevant bio-fluid using our novel ultrasensitive SMC assay. We used CSF samples from cynomolgus macaques to quantify both total HTT and pS421-HTT protein. The decision to use CSF from nonhuman primates was to ensure sufficient sample volume and to perform these proof-of-concept studies in a highly-relevant species for future potential application in a clinical setting. We detected phosphorylation at S421 in CSF from 5 different individual macaques, and the same trend as for pS421 was found in the 5 samples assessing total HTT levels, suggesting that the phosphorylated protein species detected in this matrix could be attributed to the HTT protein (Fig. 6A). Moreover, the 2B7-7D9 SMC assay was shown to specifically detect a phosphorylation signal in CSF, since dephosphorylation resulted in pS421 signals below the limit of detection, with total HTT levels virtually unaffected as a consequence of the dephosphorylation treatment (Fig. 6B). The concentration of pS421-HTT and total HTT was calculated using recombinant full length Q48 HTT, where the site of phosphorylation was confirmed by LC-MS/MS (Fig. S5 and S6) and the portion of unphosphorylated S421 could be considered negligible (Fig. S7). These data show, for the first time, that pS421-HTT could be detected in CSF from cynomolgus macaques, thereby demonstrating the ability of our SMC assay to specifically identify this critical phosphorylation event and its potential applicability in a clinical setting.

**Figure 6.**
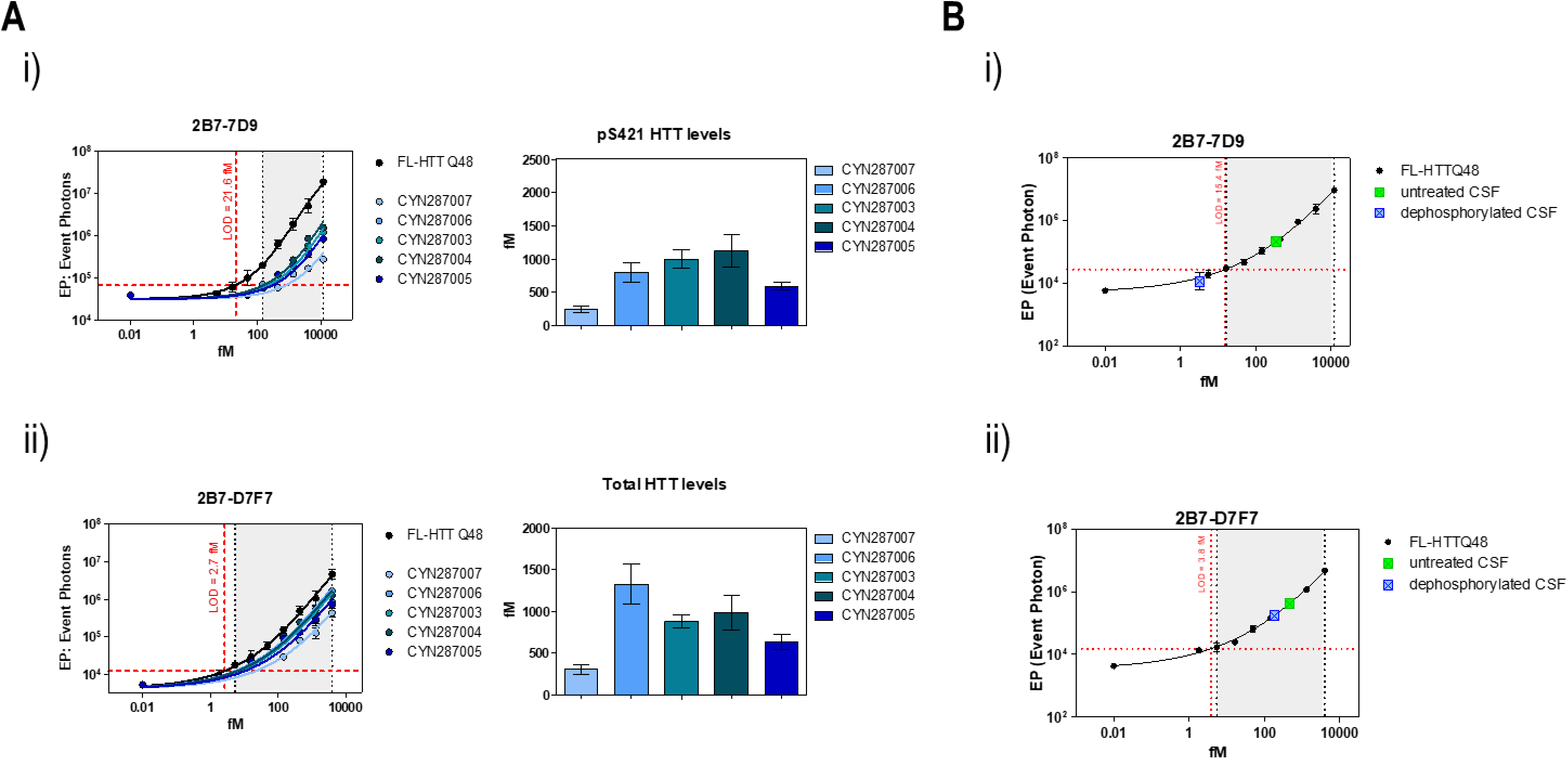
2B7-7D9 SMC assay is able to specifically detect pS421-HTT in CSF from cynomolgus macaque. **A. Comparison between pS421 and total HTT levels measured in 5 individual CSF samples from cynomolgus macaque.** i) Samples were tested in serial dilution for pS421-HTT levels (2B7-7D9 SMC assay) and for each sample the concentration of pS421-HTT (fM) was derived from the backcalculation on full-length Q48 HTT protein. ii) Same analysis in i) for total HTT levels (2B7-D7F7 SMC assay), showing comparable levels of pS421 and total HTT as assessed in the 5 CSF samples. **B. SMC analysis for pS421 and total HTT levels of untreated/treated CSF from cynomolgus macaque with lambda phosphatase.** i) pS421-HTT backcalculation of untreated or dephosphorylated CSF samples on recombinant full-length Q48 HTT protein using the 2B7-7D9 SMC assay. ii) Same analysis in i) backcalculation HTT levels employing the 2B7-D7F7 SMC assay.

Phospho-Tau at different residues is all the rage in AD and I would include reference to these studies as a paradigm for measuring phosphor-proteins in neuroD.

This is begging for testing in human case control CSF?

## Discussion

PTM status of proteins is a sensitive readout of structure-function and potential pathophysiological cell states. In prior studies we demonstrated an association of pT3 HTT and HTT CAG length, phenotypic progress in an HD mouse model and HD progression in human [25, 26]. Here we have developed a new sensitive SMC assay and demonstrate an association between pS421-HTT and HTT CAG length and phenotypic progress in an HD mouse model.

Alterations in kinase signaling and aberrant activity of specific kinases have been demonstrated in several cell and mouse models of HD, as well as in the human HD brain (reviewed in [49]). By screening a collection of kinases in HEK293T cells we demonstrate that PRKACA, the cAMP-regulated catalytic α-subunit of PKA [50], as a candidate kinase for pS421. Interestingly, progressive lowering of cAMP levels has been observed in mouse models and in post-mortem HD tissue [51], consistent with decreased PKA activity in HD that could explain the reduced levels of pS421-HTT observed by us and others [24]. The cyclic nucleotide phosphodiesterase 10A (PDE10A) is highly expressed in striatal spiny projection neurons, apparently playing a critical role in the regulation of both cAMP and cGMP signaling cascades. It has been reported that PDE10 inhibitors have a beneficial effect in HD models [52], which suggests the critical role of cAMP signalling in HD. Moreover, in the current study we have identified PRKACA as a potential kinase that can phosphorylate S421-HTT, implying that restoring cAMP levels could modulate pS421-HTT phosphorylation. Although the physiological relevance of PRKACA regulating pS421-HTT in vivo requires further investigation, our assay represents a useful tool to investigate candidate phosphorylation pathways and the functional impact of pS421-HTT modulation in HD models and clinical biomaterials.

Core CSF disease biomarkers are actively being characterized in a variety of neurodegenerative disorders, including Alzheimer’s disease, Parkinson’s disease and amyotrophic lateral sclerosis [9, 53–55]. The quantification of a number of phospho-tau sites is an exciting area of biomarker studies in AD [56, 57]. Despite the obvious relevance of CSF as a potential source of biomarkers in neurological diseases, the high variability and low protein abundance is a significant hurdle to discovering specific protein biomarkers in HD. Data in human HD case and control studies show the relevance of detecting mutant and total HTT in CSF [9, 32]. Assessment of phosphorylated proteins related to HD pathology has remained challenging due to their potential transient character together with assumed low levels in CSF, dictating the need for a highly sensitive assay. HTT phosphorylation has to date been studied in primary mouse neurons, mouse tissue, human cells and post-mortem human brain [14–16, 19, 24–29, 41, 44, 47, 58–60], but measurements have not been reported in primate CSF. To this end, we deployed our SMC assay to detect pS421 in nonhuman primate (cynomologus macaque) CSF. We were able to readily detect measurable amounts of pS421-HTT, suggesting our assay’s future utility in monitoring changes in pS421-HTT in human CSF for its potential association with HD natural history.

## Materials and Methods

### Antibodies

Monoclonal antibodies 2B7 and MW1 (which bind to the N17 domain and the polyQ stretch of HTT, respectively) were obtained from the Coriell Institute for Biomedical research (ID CH01103, ID CH00977, respectively) and their use in SMC assays has been described previously [25, 26]. The anti-pS421-HTT antibody (Ab2174, Abcam) is an affinity purified, rabbit polyclonal antibody specific for the phosphorylated S421 residue of HTT, raised against the synthetic peptide RSRSGpSIVE, corresponding to amino acids 416-424 of human HTT. Rabbit anti-pS421-HTT mAb7D9 was generated by Abclonal Technology according to standard procedures [61], using the peptide RSGpSIVELIAG comprising the HTT epitope of interest, and its characterization and specificity validation is reported in this paper. Anti-pT3 (CHDI-90001528) and anti-pS13 (CHDI-90001039) antibodies were developed by the CHDI foundation and obtained from the Coriell Institute for Biomedical research (ID CH01557 and ID CH01115) and their characterization have been reported earlier [26]. Antibody MAB2166 (Merck Millipore, catalog #MAB2166), D7F7 (Cell Signaling Technologies, catalog #5656), anti-GAPDH (Sigma-Aldrich, catalog #G9545), anti-FLAG (Sigma-Aldrich, catalog #F1804), anti-pPKA alpha (S338) (Invitrogen, catalog #PA5-64489), anti-CREB (48H2) (Cell Signaling Technologies, catalog #9197S) and anti-pCREB (Ser133) (Cell Signaling Technologies, catalog #9198S) are commercially available and used according to datasheet recommendations. Secondary antibodies used for Western blotting were Goat-anti-mouse IgG HRP conjugated (Merck Millipore, catalog #12-349) and Goat-anti-rabbit IgG HRP conjugated (Merck Millipore, catalog #12-348).

### Plasmids and proteins

The pCMV6 expression plasmids containing the cDNA coding for C-term Myc-FLAG-tagged dopamine D2 receptor (D2R; catalog #RC213065) and Protein kinase A catalytic subunit (PKA C_α_) (PRKACA; catalog #RC210332) were obtained from ORIGENE (Rockville, Maryland, United States). The cDNA encoding C-term FLAG-tagged kinase-dead PRKACA K72H/G186V (CHDI-90003842) and N-terminal HTT fragments N571Q55 with and without the S421A mutation (CHDI-90002045 and CHDI 90003841) were synthesized and subcloned into the pCDNA3.1 vector by GenScript (Nanjing, China). The cDNAs encoding full length Q23 and Q48 HTT protein with or without mutation S421A were provided by CHDI (CHDI-90002097, CHDI-90002092, CHDI-90002096 and CHDI-90002093). Expression of all constructs in HEK293T cells was validated by transient transfection as previously reported [62]. Purified and recombinant full length Q23 HTT proteins with or without S421A and D mutations were provided by CHDI (CHDI-90001858, CHDI-90002147, CHDI-90002970) as well as the recombinant full length Q48 HTT protein (CHDI-90002137, CHDI-90002962, CHDI-90002963).

### Cell culture and treatments

HEK293T cells were cultured and transiently transfected as previously described [62]. Cells were harvested 24 hours post-transfection and lysed in lysis buffer (TBS, 0.4% Triton X100) supplemented with 1X protease inhibitor cocktail (Roche, catalog #11697498001) and 1X phosphatase inhibitors (Roche, catalog #04986837001). The D2R agonist Bromocriptine (Cychem, Inc.), D2R antagonists Haloperidol (Selleck, Inc.) and Raclopride (TCI Chemicals) were delivered to HEK293T cells at a concentration of 20 µM, employing the same conditions previously reported [27]. Cell lysis was performed after 24 hours as described above. Purified, recombinant full length Q23 and Q48 HTT proteins (1 µg), overexpressed full length HTT proteins with/without S421A mutation (400 µg of total lysate) and CSF samples from cynomolgus macaque were dephosphorylated using Lambda Phosphatase (New England BioLabs, catalog #P0753L): treatments were performed with 400 U enzyme and carried out overnight (16 hrs) at 30°C as recommended by the manufacturer.

### Animal samples

CSF from cynomolgus macaque was sourced commercially (BioIVT). Tissues from *Hdh* knock-in mice (Q7/Q7 and Q7/Q175 [63]) with humanized exon 1 were obtained from the CHDI Foundation. Mouse tissue samples were homogenized using a Fast-Prep 96 (BioMedical) in 10 vol (wt/vol) of lysis buffer (TBS 0.4 % TritonX100 supplemented with a protease and phosphatase inhibitor cocktail (cOmplete Protease Inhibitor Tablets catalog #11697498001; Roche; PhosSTOP catalog #04986837001; Roche) using Lysing Matrix tubes D (MP Bio, cat#6913-500), and three lysis cycles of 30’’ at 6,000 × g were carried out. Homogenates were centrifuged and the supernatants stored at −80 °C. The quantification of lysates was performed using the BCA protein assay kit (Thermo Fisher Scientific, catalog #A53225). All mice were treated in accordance with the Guide for the Animal Care and Use of Laboratory Animals (National Research Council), and all procedures were approved by the Institutional Animal Care and Use Committee at PsychoGenics, Inc.

### Western Blotting

Samples were denatured at 95 °C in 4× Loading Buffer (125 mM Tris⋅HCl, pH 6.8, 6% SDS, 4 M urea, 4 mM EDTA, 30% glycerol, 4% 2-mercaptoethanol and bromophenol blue) and loaded on NuPAGE 4–12% Bis–Tris Gel (Thermo Fisher Scientific, catalog #WG1402BOX). Proteins were transferred on PVDF membrane (Bio-Rad Laboratories, catalog #162–0177) using wet blotting (30 V overnight at 4°C). After fixing in 0.4% paraformaldehyde solution and blocking with 5% nonfat milk in TBS/0.1% Tween-20, primary antibody incubation (final concentration: 1µg/mL in blocking solution) was carried out overnight at 4 °C, followed by secondary antibody incubations for 1 h at room temperature. Protein bands were detected using chemiluminescence substrate (Supersignal West Femto Maximum catalog #3406; Supersignal West Pico Maximum catalog #34087; Thermo Fisher Scientific) on Chemidoc XRS+ (Bio-Rad Laboratories).

### Immunoprecipitation

Immunoprecipitation was performed using Dynabeads® Protein G (Thermo Fisher Scientific, catalog #10004D) following the manufacturer’s instructions and using D7F7 and 7D9 antibody or a mock antibody (Anti-Glial Fibrillary Acidic Protein GFAP) as previously described [64]. The pulled down material was loaded on a SDS-PAGE and the Western Blot was performed as described above.

### SMC assays

The development and validation of SMC assays for the detection of PTMs and HTT protein have been previously reported [25, 26]. Briefly, the experimental protocol is as follows. 50 µL/well of dilution buffer (6% BSA, 0.8% Triton X-100, 750 mM NaCl) supplemented with protease inhibitor cocktail were added to a 96-well plate (catalog #P-96-450V-C; Axygen). Samples were diluted in artificial cerebral spinal fluid (ACSF: 0.3 M NaCl; 6 mM KCl; 2.8 mM CaCl2-2H2O; 1.6 mM MgCl2-6H20; 1.6 mM Na2HPO4-7H2O; 0.4 mM NaH2PO4-H2O) supplemented with 1% Tween-20 and protease inhibitor cocktail, in a final volume of 150 µL/well. For the capturing step, 100 µL/well of the anti-HTT antibody (MW1 or 2B7) coupled with magnetic particles diluted in Erenna Assay buffer (Merck. catalog #02-0474-00) at a final concentration of 0.025 mg/mL, was added to the assay plate, and incubated for 1 hour at room temperature under orbital shaking. The beads were then washed with Erenna System buffer (Merck, catalog #02-0111-00) and resuspended using 20 µL/well of the specific detection antibody labelled with Alexa-647 fluorophore diluted in Erenna Assay buffer at a final concentration of 0.5 ng/µl (anti-HTT pS421 Abcam #ab2174 antibody or 7D9 antibody for pS421-HTT levels readout; anti-HTT pT3 CHDI-90001528 antibody for pT3 HTT levels readout; anti-HTT pS13 CHDI-90001039 antibody for pS13 HTT levels readout; 2B7, MAB2166 (Merck Millipore) or D7F7 (Cell Signaling Technologies) for total HTT levels readout. The plate was incubated for 1 hr at room temperature under shaking. After washing, the beads were resuspended and transferred into a new 96-well plate. 10 µL/well of Erenna buffer B (Merck, catalog #02-0297-00) were added to the beads for elution and incubated for 5 min at room temperature under orbital shaking. The eluted complex was magnetically separated from the beads and transferred into a 384-well plate (Sigma, catalog #264573) where it was neutralized with 10 µL/well of Erenna buffer D (Merck, catalog #02-0368-00). Finally, the 384-well plate was heat-sealed and analyzed with the Erenna Immunoassay System.

### SMC data analysis and statistics

Relative phosphorylation signal measured by SMC assay for pS421, pT3 or pS13 was normalized on total HTT levels as the ratio between the fold increase obtained from MW1-Ab2174 or 2B7-7D9, MW1-CHDI90001528 and MW1-CHDI90001039 SMC signal (pS421, pT3, pS13 HTT levels readouts, respectively) and the fold increase derived from the 2B7-D7F7 or MW1-2B7 SMC signal (total HTT levels readouts). SMC assays for phosphorylation and total HTT levels were performed in parallel using a serial dilution (6 dilution points 1:3 plus blank, technical duplicates) of the same samples. For each readout, a curve fitting (described by a four parameter logistic curve fit) was calculated, and a fold increase between curves (which shared the top, the bottom and the slope) was assessed based on the EC50 parameter (setting one of the analyzed samples as reference). The statistical significance was assessed using a paired *t*-test (two-tailed) in the case of the analysis of n=2 groups of samples or using a One-way analysis of variance where n > 2 groups of samples were to be compared (Tukey’s Multiple Comparison Test and Dunnett’s Multiple Comparison Test). Graphs were generated and statistical analysis was performed using the software GraphPad Prism 6.

### *In vitro* kinase S421 HTT phosphorylation screening

*In vitro* screening was conducted at Reaction Biology Corporation (RBC Pennsylvania, US). A selection of 370 full length kinases were profiled for *in vitro* phosphorylation activity on the S421 epitope-derived peptide. Kinase specific control substrates were used to define 100% phosphorylation. Kinases were identified as hits when they resulted in >20% of phosphorylation of the peptides containing the S421 residue and absence of phosphorylation of the negative peptides.

### LC-MS/MS determination of S421-HTT phosphorylation

The LC-MS/MS method to assess the phosphorylation state was developed and conducted at PharmaCadence Analytical Services LLC. Purified recombinant full length HTT proteins, CHDI-90001858-34 and CHDI-90002137-20 (10 – 20 ug), were diluted with 50 mM ammonium bicarbonate and digested with trypsin/Lys C mix (1:5) for 4 hours at 37°C. LC-MS/MS analysis of the digests was conducted on a Sciex QTrap 6500+ interfaced to an eksigent express LC-200 microUPLC system. Tryptic peptides were separated by gradient elution from a 50 x 1 mm id Ace 3C18-300 column (MAC-Mod, Chadds Ford, PA) using water and acetonitrile containing 0.1% formic acid. Product ion spectra of the resultant tryptic pS421 peptide, SGpSIVELIAGGGSSCSPVLSR, were obtained and compared to that of a synthetic peptide standard at a concentration of 200 nM. Peptide fragmentation patterns were annotated using Skyline (Univ of Washington) and Protein Prospector (UCSF).

## Supplementary figures

**Figure S1 Specificity validation of 7D9 through dephosphorylation treatment of HTT protein A. Treatment by lambda phosphatase (□P) of recombinant full-length Q23 and Q48 HTT proteins.** i) WB probed with D7F7 and 7D9 antibodies revealed total HTT levels or pS421-HTT, respectively. ii) Normalized pS421 signal (2B7-7D9 SMC assay) on total HTT levels (2B7-D7F7 SMC assay) performed on samples in i). Representative experiment of n=3. **B. Treatment by lambda phosphatase (□P) of endogenous HTT expressed in HEK293T cells and overexpressing full-length HTT proteins with/without S421A mutation.** i) WB probed with D7F7 and 7D9 antibodies to demonstrate total HTT levels or pS421 HTT, respectively. Anti-GAPDH used as loading control. ii) Normalized pS421 signal (2B7-7D9 SMC assay) on total HTT levels (2B7-D7F7 SMC assay) performed on samples in i). Representative experiment of n=3.

**Figure S2 Profiling of full-length kinases for pS421 *in vitro* A.** Peptide sequence to profile 370 full length kinases for specific phosphorylation of HTT S421 *in vitro.* The consensus RXRXXS phosphorylation site is indicated. **B.** The activity of the top 10 kinases on the 4 peptides indicated in A is reported as the percentage of phosphorylation of each peptide relatively to the phosphorylation of the respective control substrate (fixed as 100%), which is established for each kinase.

**Figure S3 Profiling of full-length kinases for pS421 in HEK293T cells** Representative WB analysis of HEK293T cells transfected with the RBC screen candidate pS421-HTT kinase hits, probed with 7D9 and MAB2166 antibodies for endogenous pS421-HTT and total HTT levels, respectively. Anti-GAPDH used as loading control.

**Figure S4 Selectivity for pS421 HTT modulation by PRKACA vs other HTT phosphor-epitopes A. WB of lysates from HEK293T cells transfected with an expression plasmid for FLAG-tagged PRKACA or its catalytically inactive mutant (KD).** WB probed with CHDI-90001039 and CHDI-90001528 antibodies for pS13 and pT3 HTT levels, respectively. **B. SMC analysis of lysates from HEK293T cells transfected with an expression plasmid for FLAG-tagged PRKACA or its catalytically inactive mutant.** pS13 signal (MW1-CHDI90001039 SMC assay) and pT3 signal (MW1-CHDI90001528 SMC assay), normalized on total HTT levels (MW1-2B7 SMC assay) performed on the same samples in A. Means and standard deviations of n=3 biological replica. Statistical analysis by one-way analysis of variance (*P<0.05).

**Figure S5 LC-MS/MS on recombinant full-length HTT Q23 protein.** MS/MS spectrum of peptide SGsIVELIAGGGSSCSPVLSR produced from tryptic digestion of full length HTT Q23, CHDI-90001858-34. Predicted fragment ions observed are highlighted in green. MS/MS spectrum confirms location of phosphorylation at S421.

**Figure S6 LC-MS/MS on recombinant full-length HTT Q48 protein.** MS/MS spectrum of peptide SGsIVELIAGGGSSCSPVLSR produced from tryptic digestion of full length HTT Q48, CHDI-90002137-20. Predicted fragment ions observed are highlighted in green. MS/MS spectrum confirms location of phosphorylation at S421.

**Figure S7 Evaluation of pS421 on standard full-length HTT protein by immunoprecipitation.** WB of standard full-length HTT protein (INPUT) and standard full-length HTT protein pulled down (IP) using 7D9 or a mock antibody, probed with D7F7 and 7D9 for total and pS421-HTT levels, respectively. The blots were exposed together to allow for a direct comparison of the pS421/HTT ratio. Assuming that in the IP sample the ratio between pS421 and HTT signal equals 1, the same pS421/HTT ratio was observed for the INPUT sample.

