## Supplementary figures and images for "Modulation of huntingtin S421 phosphorylation in a Huntington’s disease mouse model and its detection in nonhuman primate cerebrospinal fluid"

### Supplementary 1

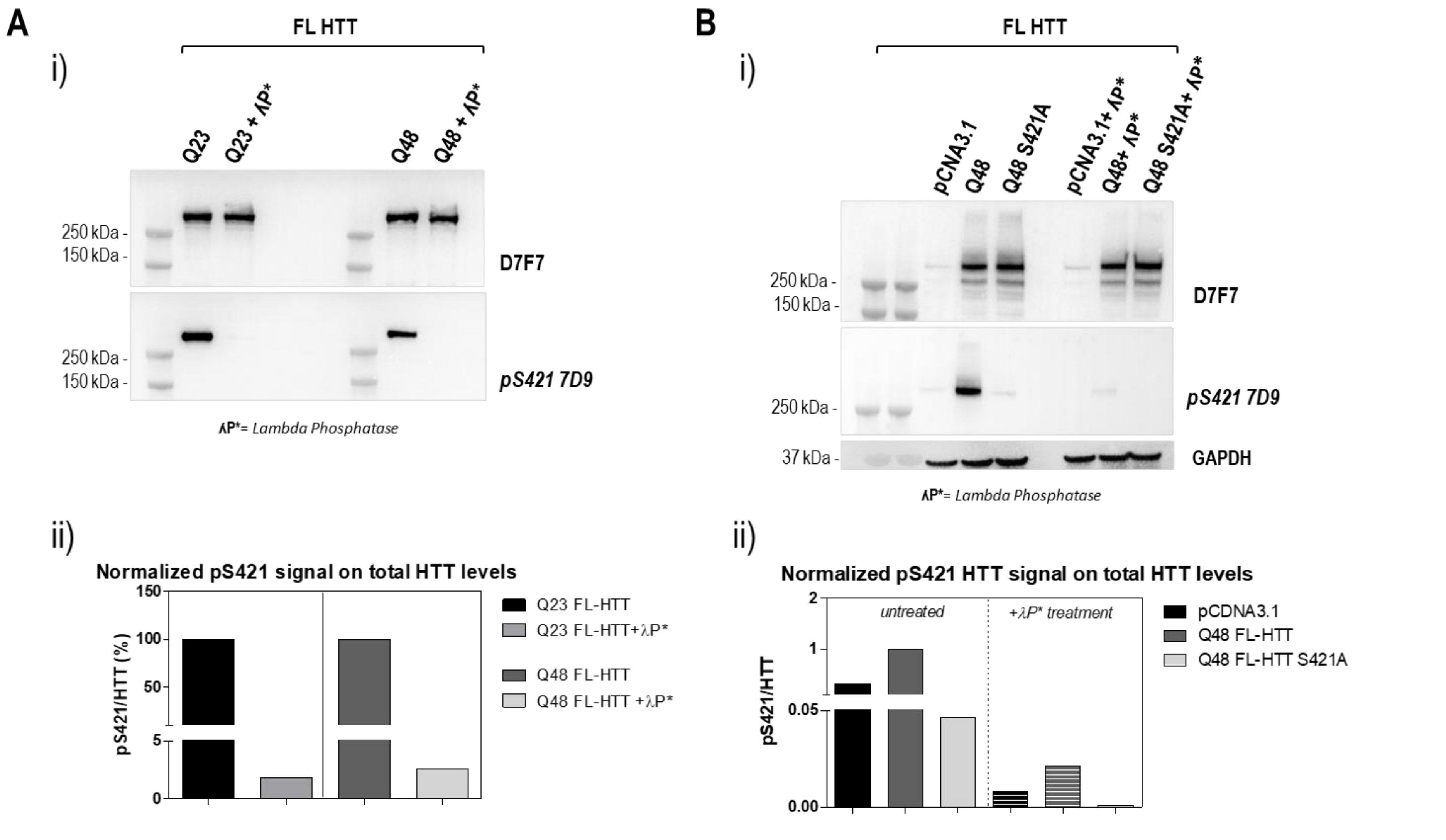

### Supplementary 2

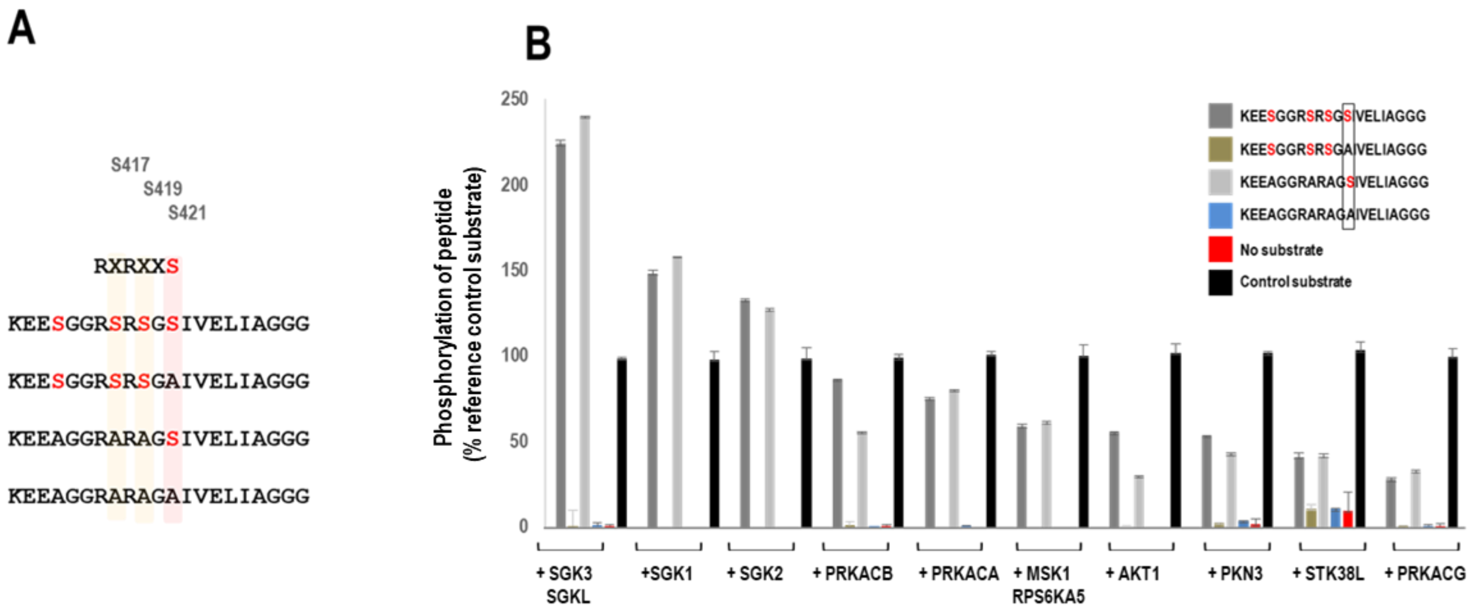

### Supplementary 3

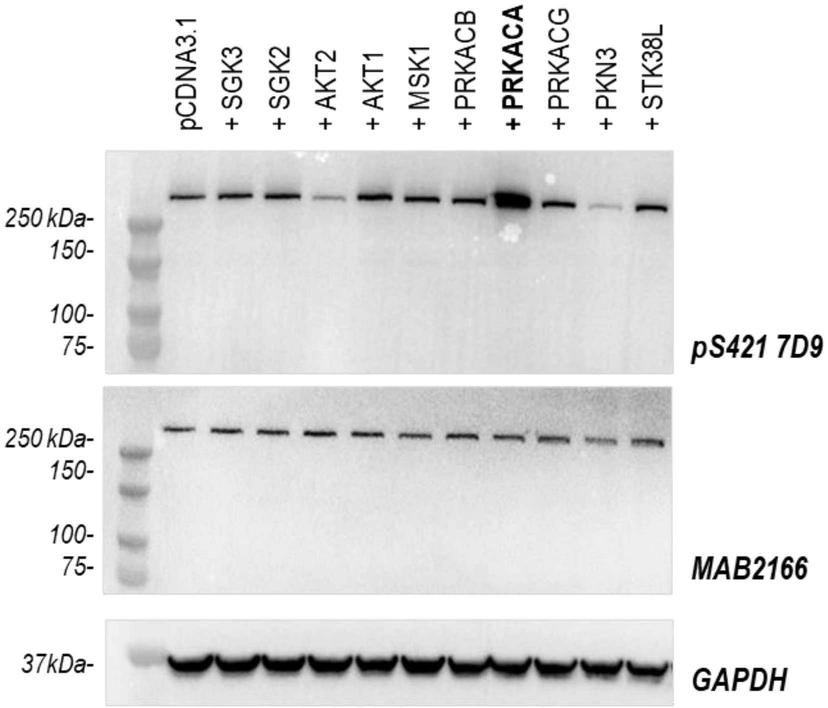

### Supplementary 4

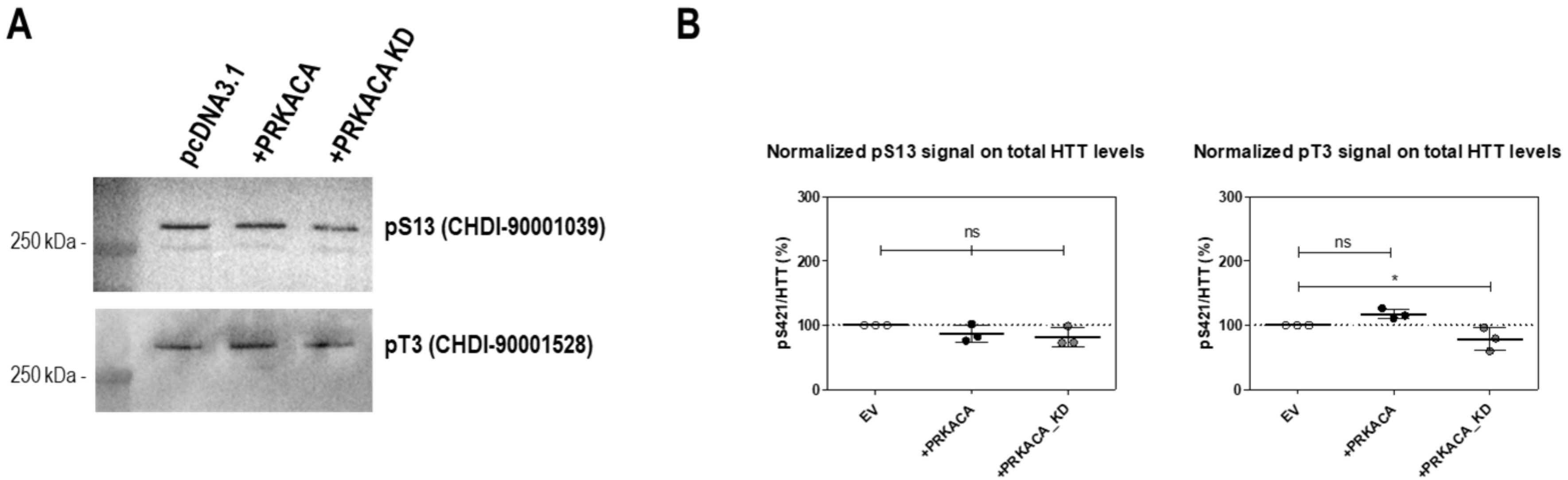

### Supplementary 5

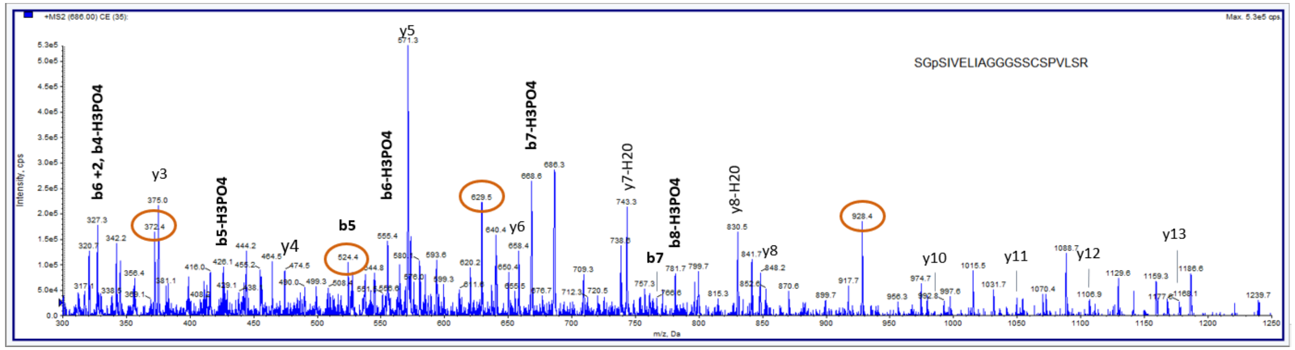


SGsIVELIAGGGSSCSPVLSR

| b-$H_2O$ | b-$H_3PO_4$ | b | | | - | y | $y^{+2}$ | y-$NH_3$ | y-$NH_3^{+2}$ | y-$H_2O$ |
|---|---|---|---|---|---|---|---|---|---|---|
| --- | --- | --- | 1 | **S** | 21 | --- | --- | --- | --- | --- |
| 127.0502 | --- | 145.0608 | 2 | **G** | 20 | 1968.946 | 984.9766 | 1951.92 | 976.4634 | 1950.936 |
| 294.0486 | 214.0822 | 312.0591 | 3 | **s** | 19 | 1911.925 | 956.4659 | 1894.898 | 947.9526 | 1893.914 |
| 407.1326 | 327.1663 | 425.1432 | 4 | **I** | 18 | 1744.926 | 872.9667 | 1727.9 | 864.4535 | 1726.916 |
| 506.201 | 426.2347 | 524.2116 | 5 | **V** | 17 | 1631.842 | 816.4247 | 1614.816 | 807.9114 | 1613.832 |
| 635.2436 | 555.2773 | 653.2542 | 6 | **E** | 16 | 1532.774 | 766.8905 | 1515.747 | 758.3772 | 1514.763 |
| 748.3277 | 668.3614 | 766.3383 | 7 | **L** | 15 | 1403.731 | 702.3692 | 1386.705 | 693.8559 | 1385.721 |
| 861.4118 | 781.4454 | 879.4223 | 8 | **I** | 14 | 1290.647 | 645.8272 | 1273.621 | 637.3139 | 1272.637 |
| 932.4489 | 852.4825 | 950.4594 | 9 | **A** | 13 | 1177.563 | 589.2851 | 1160.536 | 580.7719 | 1159.552 |
| 989.4703 | 909.504 | 1007.481 | 10 | **G** | 12 | 1106.526 | 553.7666 | 1089.499 | 545.2533 | 1088.515 |
| 1046.492 | 966.5255 | 1064.502 | 11 | **G** | 11 | 1049.504 | 525.2558 | 1032.478 | 516.7426 | 1031.494 |
| 1103.513 | 1023.547 | 1121.524 | 12 | **G** | 10 | 992.483 | 496.7451 | 975.4564 | 488.2318 | 974.4724 |
| 1190.545 | 1110.579 | 1208.556 | 13 | **S** | 9 | 935.4615 | 468.2344 | 918.4349 | 459.7211 | 917.4509 |
| 1277.577 | 1197.611 | 1295.588 | 14 | **S** | 8 | 848.4295 | 424.7184 | 831.4029 | 416.2051 | 830.4189 |
| 1380.587 | 1300.62 | 1398.597 | 15 | **C** | 7 | 761.3974 | 381.2024 | 744.3709 | 372.6891 | 743.3869 |
| 1467.619 | 1387.652 | 1485.629 | 16 | **S** | 6 | 658.3883 | 329.6978 | 641.3617 | 321.1845 | 640.3777 |
| 1564.671 | 1484.705 | 1582.682 | 17 | **P** | 5 | 571.3562 | 286.1817 | 554.3297 | 277.6685 | 553.3457 |
| 1663.74 | 1583.773 | 1681.75 | 18 | **V** | 4 | 474.3035 | 237.6554 | 457.2769 | 229.1421 | 456.2929 |
| 1776.824 | 1696.857 | 1794.834 | 19 | **L** | 3 | 375.235 | 188.1212 | 358.2085 | 179.6079 | 357.2245 |
| 1863.856 | 1783.89 | 1881.866 | 20 | **S** | 2 | 262.151 | 131.5791 | 245.1244 | 123.0659 | 244.1404 |
| --- | --- | --- | 21 | **R** | 1 | 175.119 | 88.0631 | 158.0924 | 79.5498 | --- |

### Supplementary 6

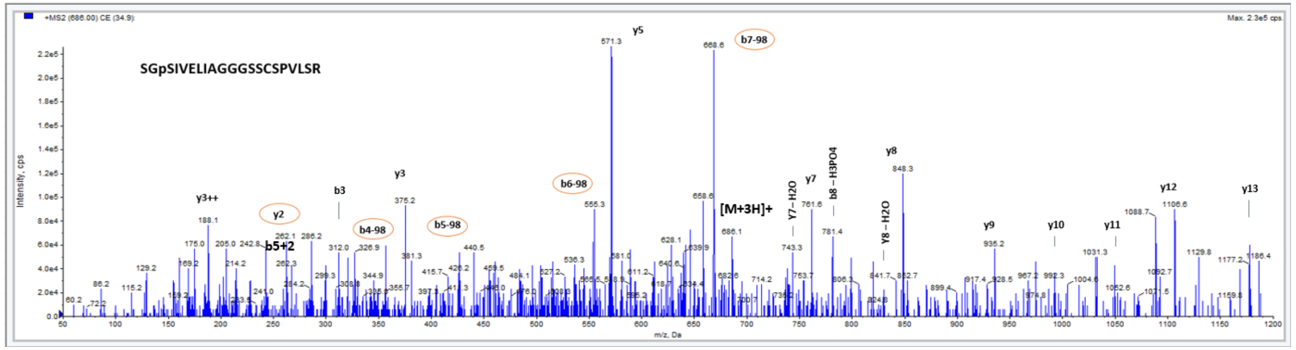


SGsIVELIAGGGSSCSPVLSR

| $b-H_2O$ | $b-H_3PO_4$ | b | | | - | y | $y^{+2}$ | $y-NH_3$ | $y-NH_3^{+2}$ | $y-H_2O$ |
|---|---|---|---|---|---|---|---|---|---|---|
| --- | --- | --- | 1 | **S** | 21 | --- | --- | --- | --- | --- |
| 127.0502 | --- | 145.0608 | 2 | **G** | 20 | 1968.946 | 984.9766 | 1951.92 | 976.4634 | 1950.936 |
| 294.0486 | 214.0822 | 312.0591 | 3 | **s** | 19 | 1911.925 | 956.4659 | 1894.898 | 947.9526 | 1893.914 |
| 407.1326 | 327.1663 | 425.1432 | 4 | **I** | 18 | 1744.926 | 872.9667 | 1727.9 | 864.4535 | 1726.916 |
| 506.201 | 426.2347 | 524.2116 | 5 | **V** | 17 | 1631.842 | 816.4247 | 1614.816 | 807.9114 | 1613.832 |
| 635.2436 | 555.2773 | 653.2542 | 6 | **E** | 16 | 1532.774 | 766.8905 | 1515.747 | 758.3772 | 1514.763 |
| 748.3277 | 668.3614 | 766.3383 | 7 | **L** | 15 | 1403.731 | 702.3692 | 1386.705 | 693.8559 | 1385.721 |
| 861.4118 | 781.4454 | 879.4223 | 8 | **I** | 14 | 1290.647 | 645.8272 | 1273.621 | 637.3139 | 1272.637 |
| 932.4489 | 852.4825 | 950.4594 | 9 | **A** | 13 | 1177.563 | 589.2851 | 1160.536 | 580.7719 | 1159.552 |
| 989.4703 | 909.504 | 1007.481 | 10 | **G** | 12 | 1106.526 | 553.7666 | 1089.499 | 545.2533 | 1088.515 |
| 1046.492 | 966.5255 | 1064.502 | 11 | **G** | 11 | 1049.504 | 525.2558 | 1032.478 | 516.7426 | 1031.494 |
| 1103.513 | 1023.547 | 1121.524 | 12 | **G** | 10 | 992.483 | 496.7451 | 975.4564 | 488.2318 | 974.4724 |
| 1190.545 | 1110.579 | 1208.556 | 13 | **S** | 9 | 935.4615 | 468.2344 | 918.4349 | 459.7211 | 917.4509 |
| 1277.577 | 1197.611 | 1295.588 | 14 | **S** | 8 | 848.4295 | 424.7184 | 831.4029 | 416.2051 | 830.4189 |
| 1380.587 | 1300.62 | 1398.597 | 15 | **C** | 7 | 761.3974 | 381.2024 | 744.3709 | 372.6891 | 743.3869 |
| 1467.619 | 1387.652 | 1485.629 | 16 | **S** | 6 | 658.3883 | 329.6978 | 641.3617 | 321.1845 | 640.3777 |
| 1564.671 | 1484.705 | 1582.682 | 17 | **P** | 5 | 571.3562 | 286.1817 | 554.3297 | 277.6685 | 553.3457 |
| 1663.74 | 1583.773 | 1681.75 | 18 | **V** | 4 | 474.3035 | 237.6554 | 457.2769 | 229.1421 | 456.2929 |
| 1776.824 | 1696.857 | 1794.834 | 19 | **L** | 3 | 375.235 | 188.1212 | 358.2085 | 179.6079 | 357.2245 |
| 1863.856 | 1783.89 | 1881.866 | 20 | **S** | 2 | 262.151 | 131.5791 | 245.1244 | 123.0659 | 244.1404 |
| --- | --- | --- | 21 | **R** | 1 | 175.119 | 88.0631 | 158.0924 | 79.5498 | --- |

### Supplementary 7

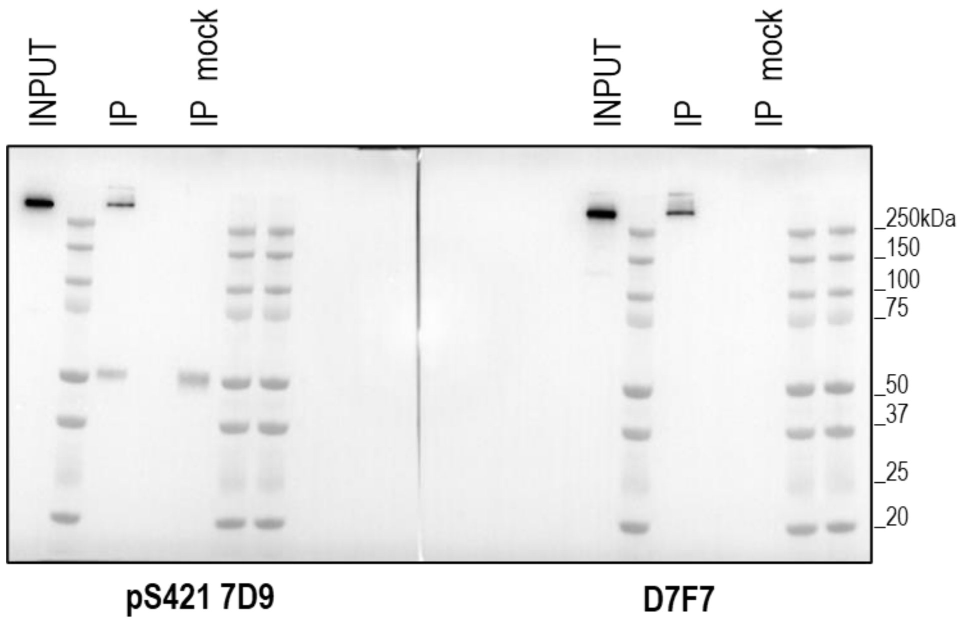
